# Comprehensive assessment of recessive, pathogenic *AARS1* alleles in a humanized yeast model reveals loss-of-function and dominant-negative effects

**DOI:** 10.1101/2024.06.20.599900

**Authors:** Molly E. Kuo, Maclaine Parish, Kira E. Jonatzke, Anthony Antonellis

## Abstract

Alanyl-tRNA synthetase 1 (*AARS1*) encodes the enzyme that ligates tRNA molecules to alanine in the cytoplasm, which is required for protein translation. Variants in *AARS1* have been implicated in early-onset, multi-system recessive phenotypes and in later-onset dominant peripheral neuropathy; to date, no single variant has been associated with both dominant and recessive diseases raising questions about shared mechanisms between the two inheritance patterns. *AARS1* variants associated with recessive disease are predicted to result in null or hypomorphic alleles and this has been demonstrated, in part, via yeast complementation assays. However, pathogenic alleles have not been assessed in a side-by-side manner to carefully scrutinize the strengths and limitations of this model system. To address this, we employed a humanized yeast model to evaluate the functional consequences of all *AARS1* missense variants reported in recessive disease. The majority of variants showed variable loss-of-function effects, ranging from no growth to significantly reduced growth. These data deem yeast a reliable model to test the functional consequences of human *AARS1* variants; however, our data indicate that this model is prone to false-negative results and is not informative for genotype-phenotype studies. We next tested missense variants associated with no growth for dominant-negative effects. Interestingly, K81T *AARS1*, a variant implicated in recessive disease, demonstrated loss-of-function and dominant-negative effects, indicating that certain *AARS1* variants may be capable of causing both dominant and recessive disease phenotypes.

## INTRODUCTION

Aminoacyl-tRNA synthetases (ARSs) are essential, ubiquitously expressed enzymes that charge tRNA molecules with cognate amino acids (Antonellis and Green, 2008). There are 37 ARS loci in the human nuclear genome, which encode enzymes that function in the cytoplasm or mitochondria (Antonellis and Green, 2008). Variants in ARS genes have been implicated in: (**1**) recessive diseases with varying clinical presentations that often include early-onset, multi-system, neurodevelopmental phenotypes; and (**2**) dominant axonal peripheral neuropathies, also called Charcot-Marie-Tooth disease (Meyer-Schuman and Antonellis, 2017). Previous genetic and functional data showed that a partial loss-of-function effect is the molecular mechanism of ARS-mediated recessive disease (Meyer-Schuman and Antonellis, 2017). Indeed, patients often carry one null and one hypomorphic alleles; complete ablation of the function of any ARS is lethal. Variants implicated in ARS-mediated dominant peripheral neuropathy affect homodimeric ARS enzymes, demonstrate loss-of-function and/or gain-of-function effects without significantly decreasing protein expression (Griffin et al., 2014; He et al., 2015), and activate the integrated stress response (Spaulding et al., 2021). We and others have presented data indicating a dominant-negative mechanism for neuropathy-associated ARS variants (Mullen et al., 2020; Meyer-Schuman et al., 2023); however, the manner in which this mechanism intersects with the integrated stress response remains unclear.

Alanyl-tRNA synthetase 1 (*AARS1*; MIM: 601065) encodes the cytoplasmic enzyme that charges tRNA^Ala^ with alanine (Antonellis and Green, 2008). Mutations in *AARS1* have been associated with multi-system recessive phenotypes and with dominant axonal peripheral neuropathy; however, no single variant has been reported to cause both dominant and recessive disease (Meyer-Schuman and Antonellis, 2017). Bi-allelic *AARS1* variants have been identified in a spectrum of recessive disease phenotypes including: (**a**) epileptic encephalopathy, hypomyelination, and progressive microcephaly; (**b**) tetraparesis; (**c**) recurrent acute liver failure; and (**d**) non-photosensitive trichothiodystrophy (**Table 1**) (Simons et al., 2015; Nakayama et al., 2017; Marten et al., 2020; Botta et al., 2021; Helman et al., 2021). The functional consequences of a subset of variants have been studied in RNA expression studies, western blot analyses, *in vitro* aminoacylation assays, and/or yeast complementation assays (**Table 2**) (Simons et al., 2015; Nakayama et al., 2017; Marten et al., 2020; Botta et al., 2021; Helman et al., 2021). However, pathogenic *AARS1* variants have not been compared in a side-by-side manner toward assessing the effectivity of each assay to detect loss-of-function effects. Addressing this issue is important for identifying a pipeline of informative functional assays that will aid in building or refuting arguments for the pathogenicity of newly identified variants.

**Table 1.**
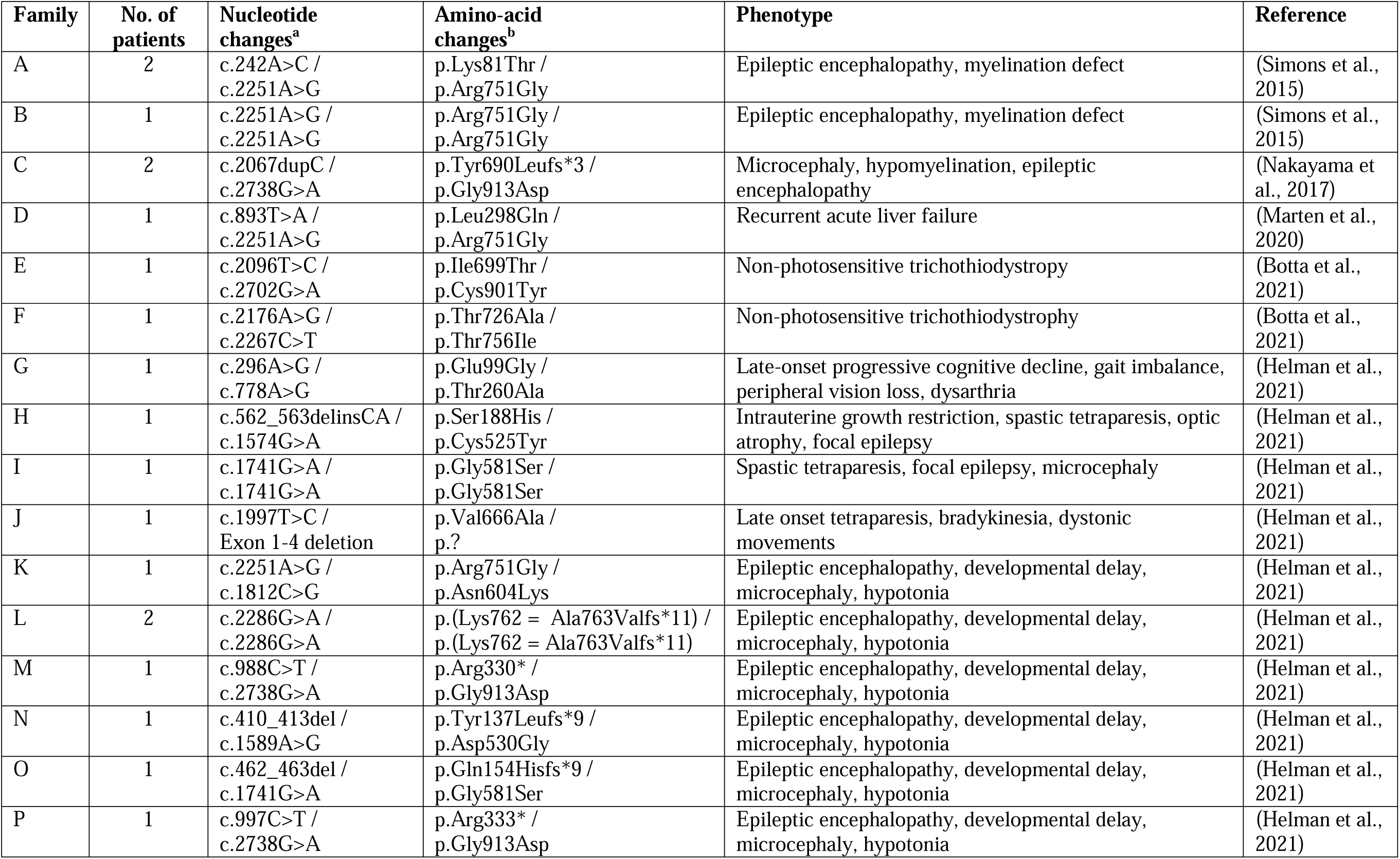
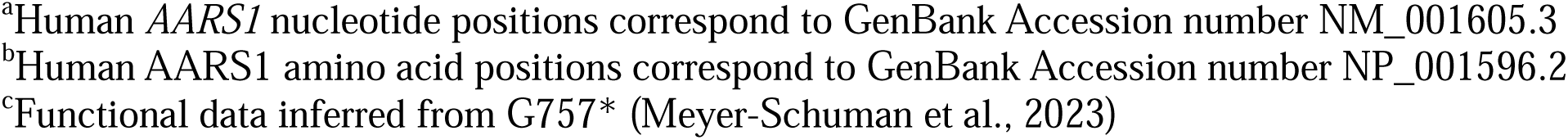
*AARS1* variants identified in patients with recessive disease phenotypes.

| Family | No. of patients | Nucleotide changes<sup>a</sup> | Amino-acid changes<sup>b</sup> | Phenotype | Reference |
| --- | --- | --- | --- | --- | --- |
| A | 2 | c.242A>C / c.2251A>G | p.Lys81Thr / p.Arg751Gly | Epileptic encephalopathy, myelination defect | (Simons et al., 2015) |
| B | 1 | c.2251A>G / c.2251A>G | p.Arg751Gly / p.Arg751Gly | Epileptic encephalopathy, myelination defect | (Simons et al., 2015) |
| C | 2 | c.2067dupC / c.2738G>A | p.Tyr690Leufs*3 / p.Gly913Asp | Microcephaly, hypomyelination, epileptic encephalopathy | (Nakayama et al., 2017) |
| D | 1 | c.893T>A / c.2251A>G | p.Leu298Gln / p.Arg751Gly | Recurrent acute liver failure | (Marten et al., 2020) |
| E | 1 | c.2096T>C / c.2702G>A | p.Ile699Thr / p.Cys901Tyr | Non-photosensitive trichothiodystrophy | (Botta et al., 2021) |
| F | 1 | c.2176A>G / c.2267C>T | p.Thr726Ala / p.Thr756Ile | Non-photosensitive trichothiodystrophy | (Botta et al., 2021) |
| G | 1 | c.296A>G / c.778A>G | p.Glu99Gly / p.Thr260Ala | Late-onset progressive cognitive decline, gait imbalance, peripheral vision loss, dysarthria | (Helman et al., 2021) |
| H | 1 | c.562_563delinsCA / c.1574G>A | p.Ser188His / p.Cys525Tyr | Intrauterine growth restriction, spastic tetraparesis, optic atrophy, focal epilepsy | (Helman et al., 2021) |
| I | 1 | c.1741G>A / c.1741G>A | p.Gly581Ser / p.Gly581Ser | Spastic tetraparesis, focal epilepsy, microcephaly | (Helman et al., 2021) |
| J | 1 | c.1997T>C / Exon 1-4 deletion | p.Val666Ala / p.? | Late onset tetraparesis, bradykinesia, dystonic movements | (Helman et al., 2021) |
| K | 1 | c.2251A>G / c.1812C>G | p.Arg751Gly / p.Asn604Lys | Epileptic encephalopathy, developmental delay, microcephaly, hypotonia | (Helman et al., 2021) |
| L | 2 | c.2286G>A / c.2286G>A | p.(Lys762 = Ala763Valfs*11) / p.(Lys762 = Ala763Valfs*11) | Epileptic encephalopathy, developmental delay, microcephaly, hypotonia | (Helman et al., 2021) |
| M | 1 | c.988C>T / c.2738G>A | p.Arg330* / p.Gly913Asp | Epileptic encephalopathy, developmental delay, microcephaly, hypotonia | (Helman et al., 2021) |
| N | 1 | c.410_413del / c.1589A>G | p.Tyr137Leufs*9 / p.Asp530Gly | Epileptic encephalopathy, developmental delay, microcephaly, hypotonia | (Helman et al., 2021) |
| O | 1 | c.462_463del / c.1741G>A | p.Gln154Hisfs*9 / p.Gly581Ser | Epileptic encephalopathy, developmental delay, microcephaly, hypotonia | (Helman et al., 2021) |
| P | 1 | c.997C>T / c.2738G>A | p.Arg333* / p.Gly913Asp | Epileptic encephalopathy, developmental delay, microcephaly, hypotonia | (Helman et al., 2021) |

<sup>a</sup>Human *AARS1* nucleotide positions correspond to GenBank Accession number NM\_001605.3
<sup>b</sup>Human *AARS1* amino acid positions correspond to GenBank Accession number NP\_001596.2
<sup>c</sup>Functional data inferred from G757\* (Meyer-Schuman et al., 2023)

**Table 2:**
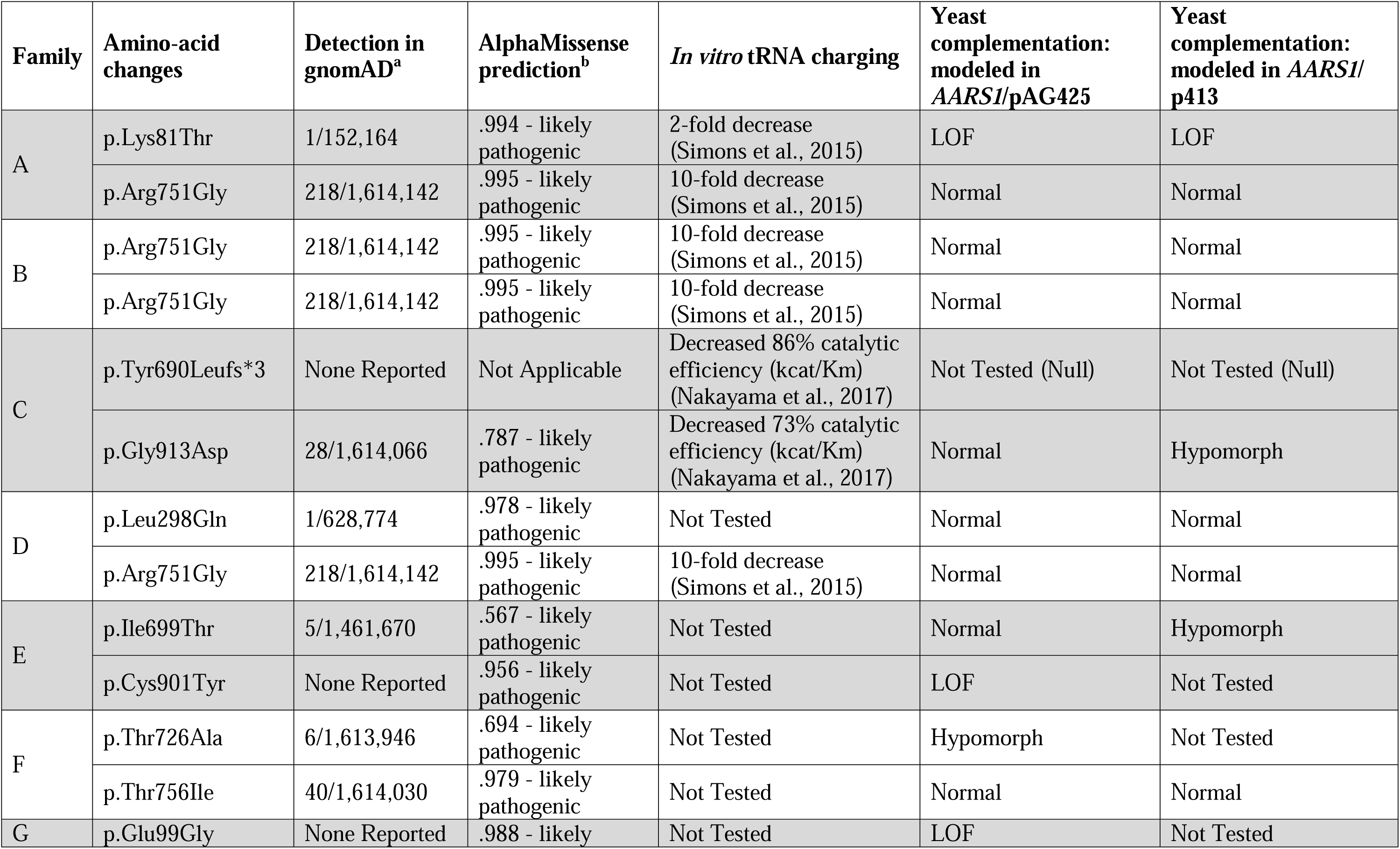
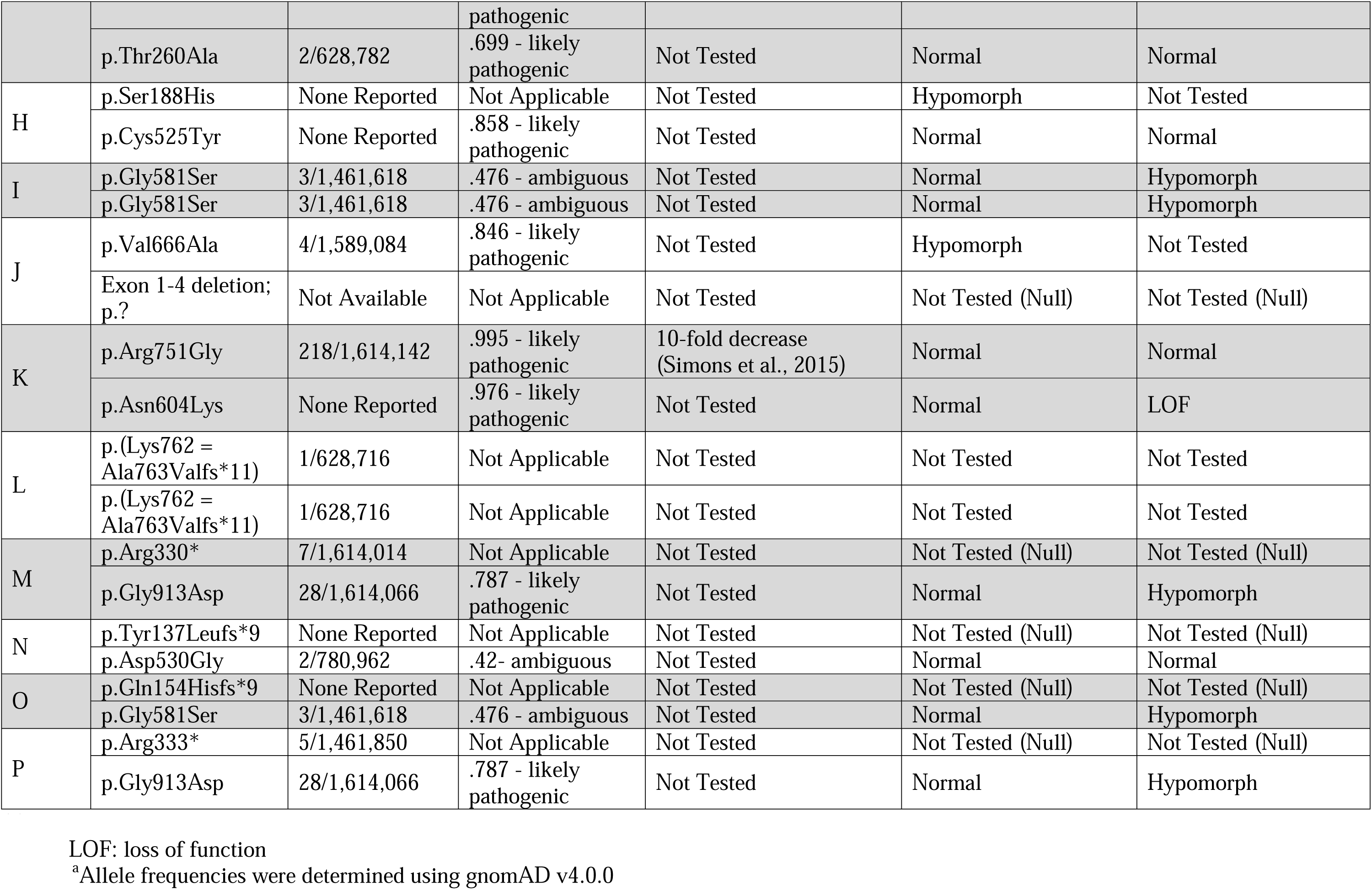
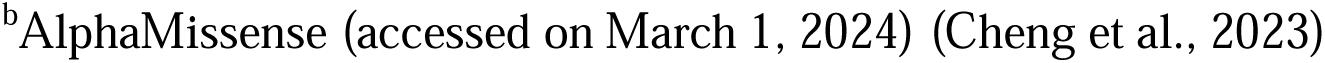
Functional consequences of disease-associated *AARS1* variants.

| Family | Amino-acid changes | Detection in gnomAD<sup>a</sup> | AlphaMissense prediction<sup>b</sup> | *In vitro* tRNA charging | Yeast complementation: modeled in *AARS1*/pAG425 | Yeast complementation: modeled in *AARS1*/p413 |
| --- | --- | --- | --- | --- | --- | --- |
| A | p.Lys81Thr | 1/152,164 | .994 - likely pathogenic | 2-fold decrease (Simons et al., 2015) | LOF | LOF |
|  | p.Arg751Gly | 218/1,614,142 | .995 - likely pathogenic | 10-fold decrease (Simons et al., 2015) | Normal | Normal |
| B | p.Arg751Gly | 218/1,614,142 | .995 - likely pathogenic | 10-fold decrease (Simons et al., 2015) | Normal | Normal |
|  | p.Arg751Gly | 218/1,614,142 | .995 - likely pathogenic | 10-fold decrease (Simons et al., 2015) | Normal | Normal |
| C | p.Tyr690Leufs*3 | None Reported | Not Applicable | Decreased 86% catalytic efficiency (kcat/Km) (Nakayama et al., 2017) | Not Tested (Null) | Not Tested (Null) |
|  | p.Gly913Asp | 28/1,614,066 | .787 - likely pathogenic | Decreased 73% catalytic efficiency (kcat/Km) (Nakayama et al., 2017) | Normal | Hypomorph |
| D | p.Leu298Gln | 1/628,774 | .978 - likely pathogenic | Not Tested | Normal | Normal |
|  | p.Arg751Gly | 218/1,614,142 | .995 - likely pathogenic | 10-fold decrease (Simons et al., 2015) | Normal | Normal |
| E | p.Ile699Thr | 5/1,461,670 | .567 - likely pathogenic | Not Tested | Normal | Hypomorph |
|  | p.Cys901Tyr | None Reported | .956 - likely pathogenic | Not Tested | LOF | Not Tested |
| F | p.Thr726Ala | 6/1,613,946 | .694 - likely pathogenic | Not Tested | Hypomorph | Not Tested |
|  | p.Thr756Ile | 40/1,614,030 | .979 - likely pathogenic | Not Tested | Normal | Normal |
| G | p.Glu99Gly | None Reported | .988 - likely pathogenic | Not Tested | LOF | Not Tested |
|  | p.Thr260Ala | 2/628,782 | .699 - likely pathogenic | Not Tested | Normal | Normal |
| H | p.Ser188His | None Reported | Not Applicable | Not Tested | Hypomorph | Not Tested |
|  | p.Cys525Tyr | None Reported | .858 - likely pathogenic | Not Tested | Normal | Normal |
| I | p.Gly581Ser | 3/1,461,618 | .476 - ambiguous | Not Tested | Normal | Hypomorph |
|  | p.Gly581Ser | 3/1,461,618 | .476 - ambiguous | Not Tested | Normal | Hypomorph |
| J | p.Val666Ala | 4/1,589,084 | .846 - likely pathogenic | Not Tested | Hypomorph | Not Tested |
|  | Exon 1-4 deletion;<br>p.? | Not Available | Not Applicable | Not Tested | Not Tested (Null) | Not Tested (Null) |
| K | p.Arg751Gly | 218/1,614,142 | .995 - likely pathogenic | 10-fold decrease (Simons et al., 2015) | Normal | Normal |
|  | p.Asn604Lys | None Reported | .976 - likely pathogenic | Not Tested | Normal | LOF |
| L | p.(Lys762 = Ala763Valfs*11) | 1/628,716 | Not Applicable | Not Tested | Not Tested | Not Tested |
|  | p.(Lys762 = Ala763Valfs*11) | 1/628,716 | Not Applicable | Not Tested | Not Tested | Not Tested |
| M | p.Arg330* | 7/1,614,014 | Not Applicable | Not Tested | Not Tested (Null) | Not Tested (Null) |
|  | p.Gly913Asp | 28/1,614,066 | .787 - likely pathogenic | Not Tested | Normal | Hypomorph |
| N | p.Tyr137Leufs*9 | None Reported | Not Applicable | Not Tested | Not Tested (Null) | Not Tested (Null) |
|  | p.Asp530Gly | 2/780,962 | .42- ambiguous | Not Tested | Normal | Normal |
| O | p.Gln154Hisfs*9 | None Reported | Not Applicable | Not Tested | Not Tested (Null) | Not Tested (Null) |
|  | p.Gly581Ser | 3/1,461,618 | .476 - ambiguous | Not Tested | Normal | Hypomorph |
| P | p.Arg333* | 5/1,461,850 | Not Applicable | Not Tested | Not Tested (Null) | Not Tested (Null) |
|  | p.Gly913Asp | 28/1,614,066 | .787 - likely pathogenic | Not Tested | Normal | Hypomorph |

LOF: loss of function
<sup>a</sup>Allele frequencies were determined using gnomAD v4.0.0
<sup>b</sup>AlphaMissense (accessed on March 1, 2024) (Cheng et al., 2023)

In this study, we employ a humanized yeast model to comprehensively evaluate the functional consequences of all 16 reported *AARS1* missense variants associated with recessive disease; we assume that all identified early frameshift, nonsense, and large-deletion mutations represent null alleles. Each missense variant was individually tested in one or two yeast complementation assays, which employ high- or low-copy number expression vectors. Subsequently, loss-of-function missense variants were evaluated for dominant-negative effects by testing the ability of each to repress a wild-type copy of human *AARS1*. Our data revealed that yeast is an informative model to test human *AARS1* variants for loss-of-function effects in the context of recessive disease. These data also revealed important limitations of yeast as a model system including an inability to: (***i***) detect subtle deficits in gene function; and (***ii***) explain genotype-phenotype relationships. It will be important to consider these limitations in future uses of yeast to study pathogenic *AARS1* alleles. In addition, our data suggest that all loss-of-function *AARS1* missense variants have the potential to exert dominant-negative effects, but that for some variants this is ameliorated through, for example, reduced protein expression or dimerization with the wild-type subunit. These findings suggest that some carriers of pathogenic *AARS1* alleles that cause recessive disease (*i.e.*, the parents and siblings of affected individuals) may manifest *AARS1*-associated dominant axonal neuropathy. In sum, our findings have important implications for studying the allelic and clinical heterogeneities—as well as the mechanisms—of *AARS1*-associate inherited disease.

## MATERIALS AND METHODS

### Allele frequencies and conservation

The frequency of each variant was collected from gnomAD v4.0.0 (Lek et al., 2016; Karczewski et al., 2020). Conservation of each variant was examined by aligning AARS1 protein orthologs from multiple species with Clustal Omega (https://www.ebi.ac.uk/Tools/msa/clustalo/). The accession numbers used were: human (*Homo sapiens*, NP_001596.2), mouse (*Mus musculus*, NP_666329.2), zebrafish (*Danio rerio*, NP_001037775.1), fly (*Drosophila melanogaster*, AAF05593.1), worm (*Caenorhabditis elegans*, O01541.1), yeast (*Saccharomyces cerevisiae*, EDN63655.1), and bacteria (*Escherichia coli*, BAA16559.1).

### Yeast complementation assays

The 16 *AARS1* missense variants studied were modeled in the human *AARS1* open-reading frame. We performed site-directed mutagenesis using a sequence-verified, wild-type *AARS1* pDONR221 construct (Invitrogen), mutagenesis primers specifically designed for each *AARS1* missense variant (primer sequences available in Supplementary Table 1), and the Quikchange II Site-Directed Mutagenesis Kit (Agilent). After transformation into bacteria, individual colonies were collected, and DNA was isolated. We next subjected each DNA sample to Sanger sequencing analysis to verify the *AARS1* variant sequence and the absence of unintended mutations that could arise from polymerase error. The sequence-verified expression constructs for wild-type and mutated *AARS1* were then cloned into either the pAG425 expression vector (pAG425GAL-ccdB; Addgene #14153; pAG425 harbors a galactose-inducible promoter) or the p413 expression vector (ATCC #87370; p413 harbors the constitutive *ADH* promoter) using Gateway cloning technology (Invitrogen).

For expression of *AARS1* alleles from a <u>high</u>-copy number vector, null (G757* *AARS1*, for which no protein product is expressed [(Meyer-Schuman et al., 2023)]), wild-type, or mutant *AARS1* in pAG425 expression constructs were individually transformed into a haploid yeast strain, ptetO7-*ALA1* (p*ALA1*::kanR-tet07-TATA *URA3*::CMV-tTA *MATa*; from the Yeast Tet-Promoters Hughes Collection, Horizon Discovery, accession: YSC1180), and subsequently plated on media lacking leucine (pAG425 harbors the *LEU2* gene). Colonies were picked into liquid media lacking leucine and grown at 30°C and shaking at 275 rpm for 48 hours. Aliquots of these samples were diluted 1:10 and 1:100 on three plates (Takara Bio): (**1**) glucose plates lacking leucine (endogenous *ALA1* is expressed and *AARS1* on pAG425 is not expressed); (**2**) galactose/raffinose plates lacking leucine (endogenous *ALA1* and *AARS1* on pAG425 are both expressed); and (**3**) galactose/raffinose plates lacking leucine and with 10 ug/ml doxycycline (endogenous *ALA1* is repressed and *AARS1* on pAG425 is expressed).

For expression of *AARS1* alleles from a <u>low</u>-copy number vector, all experimental and control alleles (see above) in p413 constructs were individually transformed into the haploid yeast strain mentioned above and subsequently plated on media lacking histidine (p413 harbors the *HIS3* gene). Colonies were picked into liquid media lacking histidine and grown at 30°C and shaking at 275 rpm for 48 hours. Aliquots of these samples were diluted 1:10 and 1:100 on two plates (Takara Bio): (**1**) glucose plates lacking histidine (endogenous *ALA1* is expressed and *AARS1* on p413 is expressed); and (**2**) glucose plates lacking histidine with 10 ug/ml doxycycline (endogenous *ALA1* is repressed and *AARS1* on p413 is expressed).

For dominant toxicity assays, null (G757* *AARS1*), wild-type, or mutant *AARS1* in pAG425 expression constructs were transformed into the ptetO7-*ALA1* haploid yeast strain along with either empty or wild-type *AARS1* p413. Transformed yeast were plated on media lacking leucine (pAG425 harbors the *LEU2* gene) and histidine (p413 harbors the *HIS3* gene), and colonies were picked into liquid media lacking leucine and histidine. Aliquots were spotted on three plates: (**1**) glucose plates lacking leucine and histidine (endogenous *ALA1* is expressed, *AARS1* on p413 is expressed, and *AARS1* on pAG425 is not expressed); (**2**) galactose/raffinose plates lacking leucine and histidine (endogenous *ALA1, AARS1* on p413, and *AARS1* on pAG425 are all expressed); and (**3**) galactose/raffinose plates lacking leucine and histidine and with 10 ug/ml doxycycline (endogenous *ALA1* is repressed, *AARS1* on p413 is expressed, and *AARS1* on pAG425 is expressed).

After plating, yeast growth and viability were assessed visually after 5 days at 30°C. At least two independent transformations were performed and at least three colonies per transformation were analyzed. Images of yeast spots were quantified to assess the relative growth of mutant variants in comparison to wild-type *AARS1* using an established protocol (Petropavlovskiy et al., 2020). Images from the control (-his or -his-leu glucose) and experimental (-his or -his-leu galactose/raffinose with doxycycline) plates were imported to ImageJ. The image background was subtracted and assessed for uniformity across the image. The density of cells in each yeast spot (1:10 dilution) was individually measured. Growth for each spot on the experimental plate was calculated relative to the corresponding spot on the control plate. Relative growth for each mutant was then normalized to the growth of the wild-type spot on that plate. The average growth rate across three colony replicates for each mutant was calculated. Statistical significance was determined using GraphPad Prism, which performed one-way ANOVA with the Geisser-Greenhouse correction and Dunnett’s multiple comparison’s test with individual variances computed for each comparison.

### Protein isolation and western blot analysis

Haploid yeast (ptetO7-*ALA1*) were transformed with wild-type, G757*, or mutant *AARS1* pAG425 constructs as above. Transformation colonies were picked into 5 ml galactose/raffinose liquid media lacking leucine and incubated at 30°C and shaking at 275 rpm for 2 days. Protein was isolated from yeast as previously described (Meyer-Schuman et al., 2023). Briefly, yeast cells were pelleted, supernatant was removed, and samples were frozen at -80°C. Samples were resuspended in 50 ul lysis buffer (50 mM Na-HEPES pH 7.5, 200 mM NaOAc, 1 mM EDTA, 0.25% NP-40, 3mM DTT, and 1X Halt Protease Inhibitor Cocktail [Thermo Fisher Scientific]). Approximately 100 ul of 0.5 mm glass beads were added to each sample, and samples were vortexed at 4°C for 3 minutes, then incubated on ice for 3 minutes, then vortexed again for 3 minutes. The bottom of each 1.5 ml tube was punctured with a 26-guage needle and the tube was placed into a 14-mL polypropylene round-bottom tube before centrifuging at 4°C for 5 minutes at 200 rcf. The lysates were collected and protein concentration was quantified using the Thermo Scientific Pierce BCA Protein Assay kit.

For western blot analyses, 50 ug per sample was analyzed. Protein samples were prepared with 1X SDS sample buffer (Thermo Fisher Scientific) and 2-mercaptoethanol. Samples were denatured at 99°C for 5 minutes and separated on a 4-20% tris-glycine protein gel (Thermo Fisher Scientific) at 150 V for 75 minutes. Protein was transferred to a methanol-treated polyvinylidene difluoride membrane (Millipore Sigma) using a Mini Trans Blot Electrophoretic Transfer Cell (Biorad) at 100 V for 1 hour. The membrane was blocked with 5% milk solution for 1 hour at room temperature and then incubated overnight at 4°C in blocking solution containing primary antibodies, anti-AARS1 (Bethyl; A303-473A; 1: 1,000) and anti-PGK1 (Abcam; ab113689; 1:3,000). The membrane was washed with 1X tris-buffered saline solution and Tween-20 (TBST) for 5 minutes three times and then incubated with blocking solution containing IRDye 800CW goat anti-rabbit IgG secondary antibody (LI-COR), IRDye 680RD goat anti-mouse IgG secondary antibody (LI-COR), 0.02% SDS, and 0.1% Tween-20 for 1 hour at room temperature. The membrane was washed in 1X TBST for 5 minutes three times and imaged on a LI-COR Odyssey CLx Imager. Each experiment was performed three times. Band intensity was measured using ImageJ and for each sample the intensity of the AARS1 band was calculated relative to the PGK1 band and then normalized to the intensity of the band for yeast transformed with the wild-type *AARS1* construct. Statistical significance was determined using GraphPad Prism, which performed one-way ANOVA with the Geisser-Greenhouse correction and Dunnett’s multiple comparison’s test with individual variances computed for each comparison.

## RESULTS

### A collection AARS1 variants currently implicated in recessive disease

To collect all reported recessive disease-associated *AARS1* alleles, we performed a literature review, which revealed 23 variants (**Table 1**) (Simons et al., 2015; Nakayama et al., 2017; Marten et al., 2020; Botta et al., 2021; Helman et al., 2021). Seven variants result in premature stop codons or large deletions, and we assume that these genetic lesions result in null alleles. This assumption is based, in part, on our previous observation that the G757* engineered *AARS1* allele results in a lack of protein expression (Meyer-Schuman et al., 2023) and that the 3’-most premature stop codon occurs at A763. In addition, 16 of the reported disease-associated variants are missense changes. The AARS1 protein contains: (**1**) an aminoacylation domain for activating and transferring the amino acid to tRNA; (**2**) a tRNA recognition domain; (**3**) an editing domain for hydrolyzing mischarged tRNA; and (**4**) a C-terminal domain that is important for dimerization (Naganuma et al., 2009). The variants localize throughout the functional domains. Of the missense variants, three are located in the aminoacylation domain, two are located in the tRNA recognition domain, nine are located in the editing domain, and two are located in the C-terminal domain (**Figure 1A**). The affected residues for each of the 16 missense variants show variable conservation, with all residues conserved among human, mouse, and fish (**Figure 1B**). Each variant was either not present or present at a low frequency (< 0.0002) in gnomAD and no homozygous individuals were noted (**Table 2**). Combined, these data are consistent with the disease-associated *AARS1* missense variants being pathogenic and causing loss-of-function effects.

**Figure 1.**
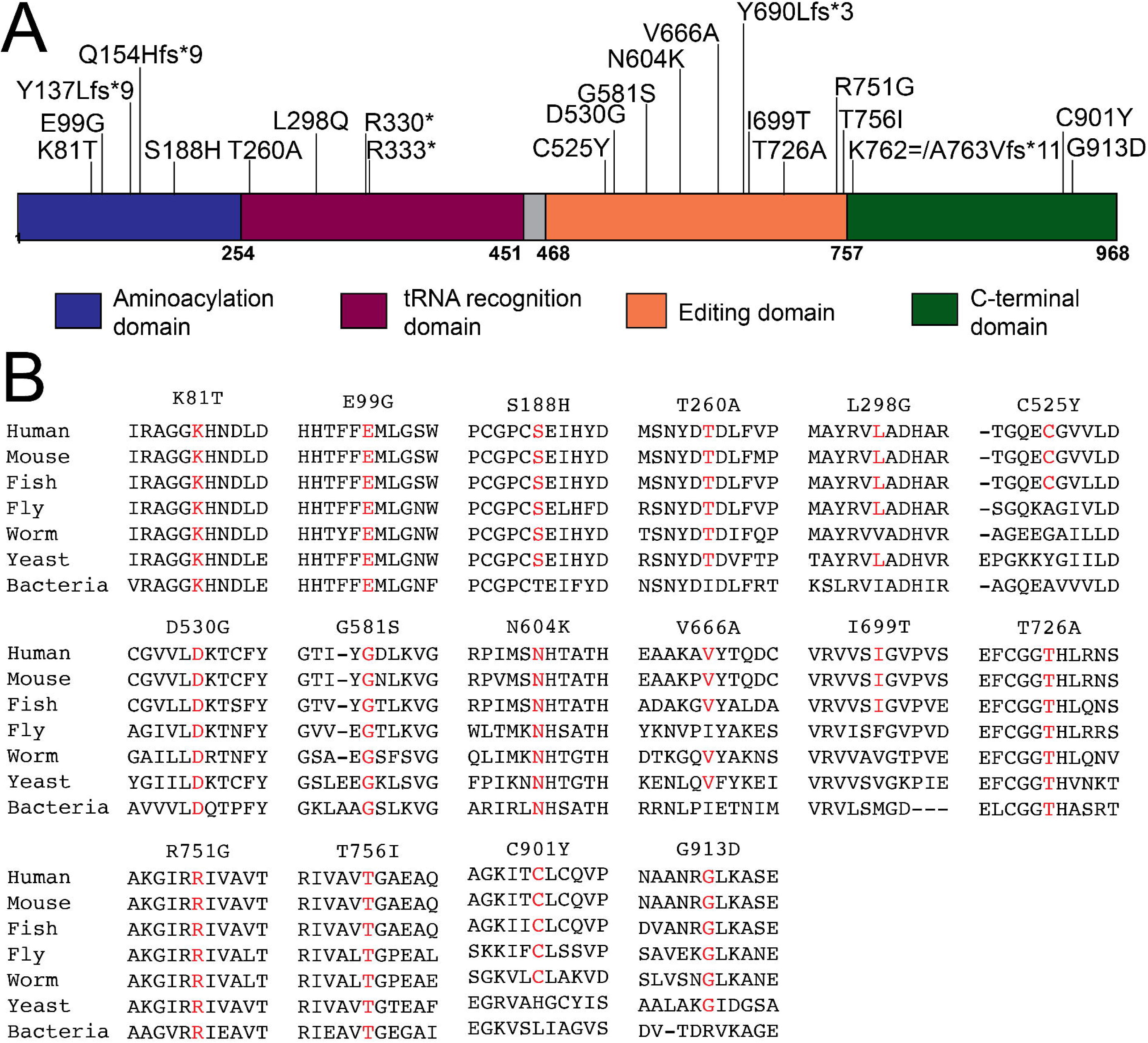
Localization and conservation of *AARS1* variants implicated in recessive disease. (**A**) AARS1 functional domains are indicated in blue (aminoacylation domain), purple (tRNA recognition domain), orange (editing domain), and green (C-terminal domain). The positions of the variants are shown across the top and numbers along the bottom indicate amino-acid positions. The exon 1-4 deletion from Family J is not depicted. (**B**) Conservation of the affected amino-acid residues for all missense variants implicated in recessive disease. The position of each variant is shown along with flanking AARS1 amino-acid residues from multiple, evolutionarily diverse species. The wild-type human amino-acid residue at the position of the affected residue is shown in red for each species.

### AARS1 variants implicated in recessive disease show loss-of-function effects in yeast complementation assays

To test and directly compare the functional consequences of the 16 disease-associated *AARS1* missense variants, we employed a humanized yeast complementation assay and tested the ability of each variant to complement loss of the endogenous yeast gene, *ALA1*. Briefly, we used a haploid yeast strain (ptetO7-*ALA1*) with endogenous yeast *ALA1* under the control of a tetracycline-repressible promoter (Meyer-Schuman et al., 2023). Wild-type *AARS1*, mutant *AARS1*, or a previously reported null allele (G757*) were cloned into the pAG425 vector, a high-copy number vector with a galactose-inducible promoter. These expression constructs were transformed into the yeast strain and yeast growth was evaluated on medium containing doxycycline (to represses endogenous *ALA1* expression) and galactose (to expresses *AARS1* from pAG425). Null *AARS1* did not support yeast growth (**Figure 2A** and **2B**, and Supplementary Figure 1), consistent with *ALA1* being an essential gene. Wild-type human *AARS1* expression supported yeast growth (**Figure 2A** and **2B**, and Supplementary Figure 1), indicating that human *AARS1* can complement the loss of the endogenous *ALA1* locus, which is consistent with previous findings (Meyer-Schuman et al., 2023). Three human missense variants (K81T, E99G, and C901Y *AARS1*) were unable to support yeast growth (**Figure 2A** and **2B**, and Supplementary Figure 1), indicating that these variants represent null alleles. Three additional human variants (S188H, V666A, and T726A *AARS1*) supported growth that was significantly reduced compared to wild-type *AARS1* (p-value < 0.05; **Figure 2A** and **2B**, and Supplementary Figure 1), indicating that these variants are hypomorphic alleles. The 10 remaining human variants (T260A, L298Q, C525Y, D530G, G581S, N604K, I699T, R751G, T756I, and G913D *AARS1*) supported growth in a manner similar to wild-type *AARS1* (**Figure 2A** and **2B**, and Supplementary Figure 1).

**Figure 2.**
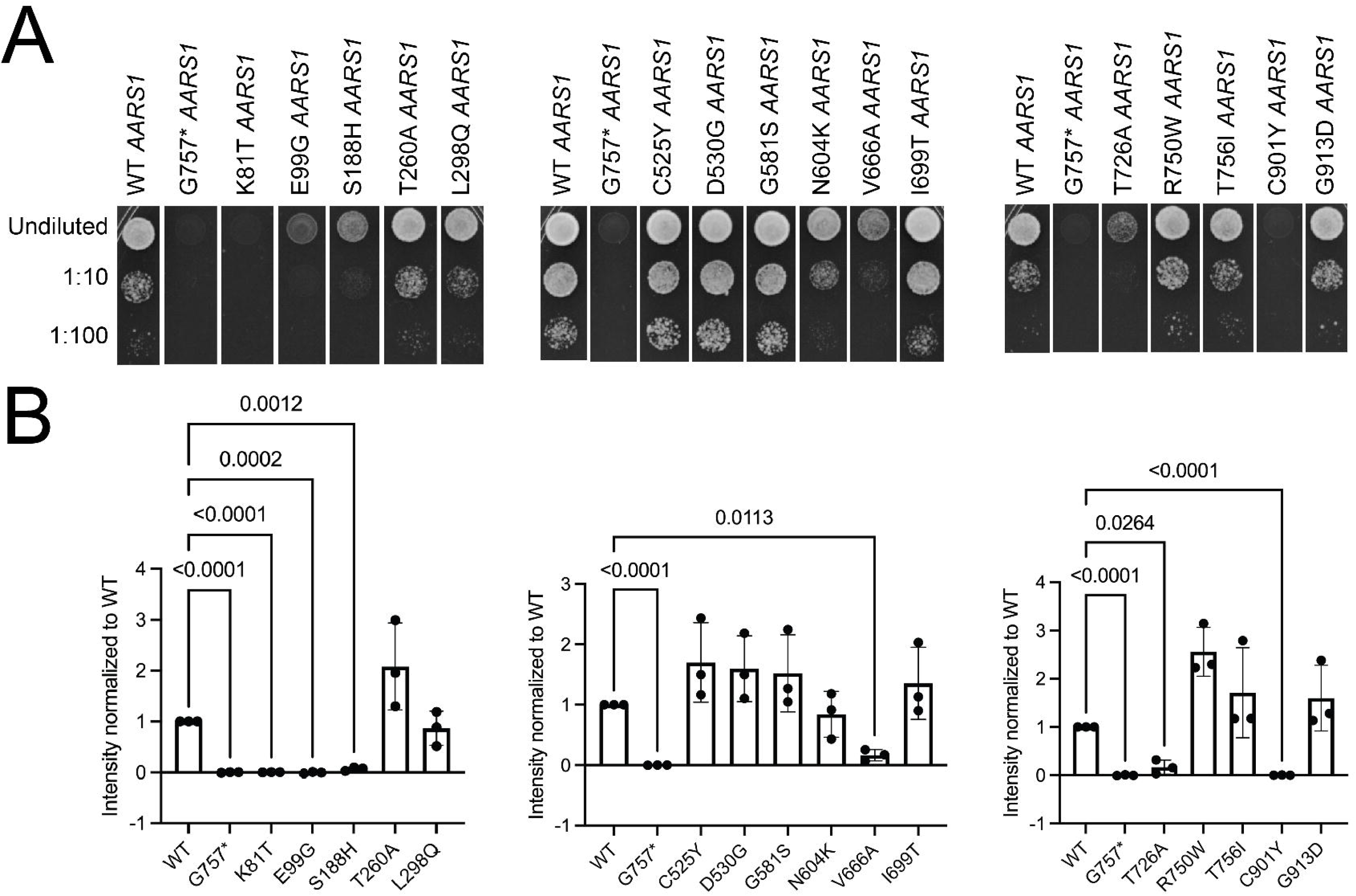
Assessment of loss-of-function effects of *AARS1* missense variants in yeast complementation assays using a high-copy number vector (pAG425). Haploid yeast with a doxycycline-repressible endogenous *ALA1* (the yeast ortholog of *AARS1*) were transformed with pAG425 vectors containing wild-type (WT) or mutant *AARS1*, or a vector with a null allele (G757* *AARS1*); the vector used in each experiment is indicated across the top. (**A**) Resulting cultures were plated undiluted or diluted (1:10 or 1:100) on media containing doxycycline and grown at 30°C for five days. (**B**) Images of yeast spots were quantified to assess the relative growth of mutant variants in comparison to wild-type *AARS1.* The average growth rate across three colony replicates for each mutant was calculated and is depicted as the bar height. Error bars represent standard deviation. Statistical significance was determined by one-way ANOVA with the Geisser-Greenhouse correction and Dunnett’s multiple comparison’s test with individual variances computed for each comparison. Only comparisons that were statistically significant are annotated with a p-value.

In the above experiments, each variant was expressed from a high-copy number plasmid, raising the possibility of false-negative results (*i.e.*, by masking subtle loss-of-function effects). For the 10 missense variants that supported similar growth to wild-type *AARS1* in the above system, we evaluated effects on yeast growth using a low-copy number expression vector. Here, wild-type, mutant, and null (G757*) human *AARS1* were cloned into p413, a low-copy number plasmid.

The p413 expression constructs were then transformed into the ptetO7-*ALA1* haploid yeast strain and growth was evaluated on doxycycline-containing medium to repress endogenous *ALA1*. Four human alleles (G581S, N604K, I699T, and G913D *AARS1*) demonstrated significantly reduced growth compared to wild-type *AARS1* (**Figure 3A** and **3B**, and Supplementary Figure 2). The six remaining human alleles (T260A, L298Q, C525Y, D530G, R751G, and T756I *AARS1*) supported growth similar to wild-type human *AARS1* (**Figure 3A** and **3B** and Supplementary Figure 2). Thus, the humanized yeast complementation assay described above was able to detect loss-of-function effects for 10 out of 16 (62.5%) disease-associated *AARS1* missense variants.

**Figure 3.**
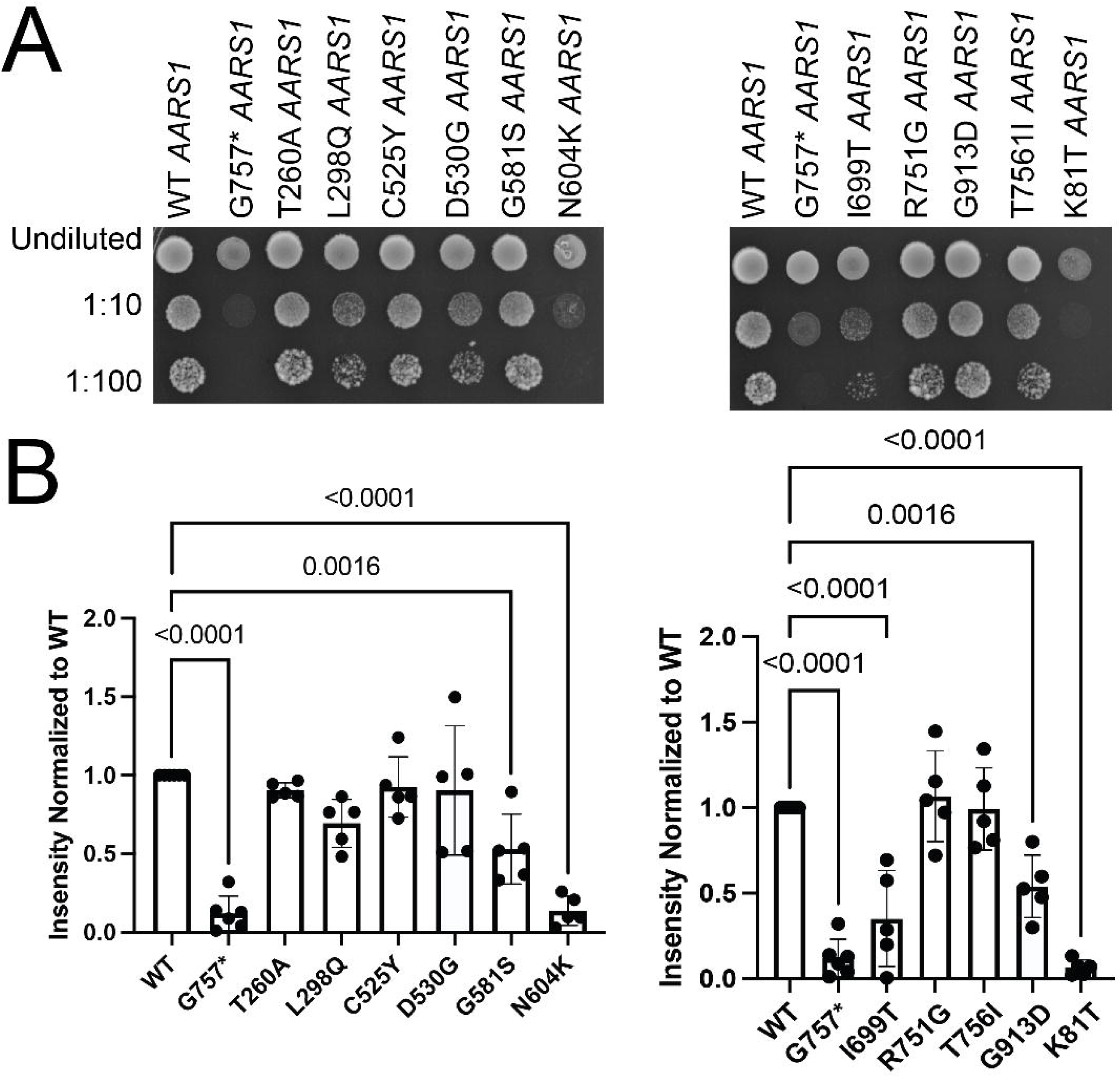
Assessment of loss-of-function effects of *AARS1* missense variants in yeast complementation assays with low-copy number vector (p413). Haploid yeast with a doxycycline-repressible endogenous *ALA1* (the yeast ortholog of *AARS1*) were transformed with p413 vectors containing wild-type (WT) or mutant *AARS1*, or a null allele (G757* *AARS1*); the vector used in each experiment is indicated across the top. (**A**) Resulting cultures were plated undiluted or diluted (1:10 or 1:100) on media containing doxycycline and grown at 30°C for five days. (**B**) Images of yeast spots were quantified to assess the relative growth of mutant variants in comparison to wild-type *AARS1.* The average growth rate across three colony replicates for each mutant was calculated and is depicted as the bar height. Error bars represent standard deviation. Statistical significance was determined by one-way ANOVA with the Geisser-Greenhouse correction and Dunnett’s multiple comparison’s test with individual variances computed for each comparison. Only comparisons that were statistically significant are annotated with a p-value.

### K81T AARS1 results in protein expression comparable to wild-type AARS1 in yeast assays

Loss-of-function effects have been described for *AARS1* alleles associated with recessive syndromes and with dominant axonal neuropathy (McLaughlin et al., 2011; Simons et al., 2015). While a single *AARS1* allele has not been implicated in both recessive and dominant phenotypes, the neuropathy is later-onset and may not have manifested at (time of examination) in the parents or siblings of individuals affected with the *AARS1*-related recessive disease. Additionally, a later-onset peripheral neuropathy may not be a primary concern of families with members affected with *AARS1*-related recessive phenotypes. Previously, we demonstrated that neuropathy-associated *AARS1* alleles have dominant-negative properties in a humanized yeast assay (Meyer-Schuman et al., 2023). To determine if certain recessive *AARS1* alleles also have dominant-negative properties, we studied the three missense *AARS1* variants that resulted in a complete loss-of-function effect: K81T, E99G, and C901Y.

For an allele to have dominant-negative properties, the genetic lesion should not disrupt expression of the gene product (Veitia, 2007). To assess this for the three human *AARS1* alleles under study, we performed western blot analyses on protein isolated from yeast transformed with pAG425 vectors to express wild-type, null, or mutant human *AARS1*. We used antibodies against AARS1 and phosphoglycerate kinase (PGK1), which was utilized as a yeast cell loading control. In protein isolated from yeast transformed with G757* *AARS1*, there was no band, consistent with G757* *AARS1* being a null allele and resulting in no protein expression (**Figure 4A** and Supplementary Figure 3). In protein isolated from yeast transformed with wild-type *AARS1*, there is a band between 100 and 130 kDa, which is consistent with the predicted molecular weight of the AARS1 protein, ∼107 kDa (**Figure 4A** and Supplementary Figure 3). In yeast transformed with K81T or E99G *AARS1*, AARS1 protein band intensity was similar to yeast transformed with wild-type *AARS1*, suggesting that these two variants do not disrupt AARS1 protein expression (**Figure 4A** and Supplementary Figure 3; note that the mean E99G protein level is decreased but not significantly different from wild-type). Finally, in protein isolated from yeast transformed with C901Y *AARS1*, AARS1 protein band intensity was significantly reduced compared to yeast transformed with wild-type *AARS1* (p-value 0.0003; **Figure 4A** and Supplementary Figure 3). In summary, these studies show that K81T and E99G AARS1 are loss-of-function proteins that are expressed at levels comparable to wild-type AARS1 in yeast cells.

**Figure 4.**
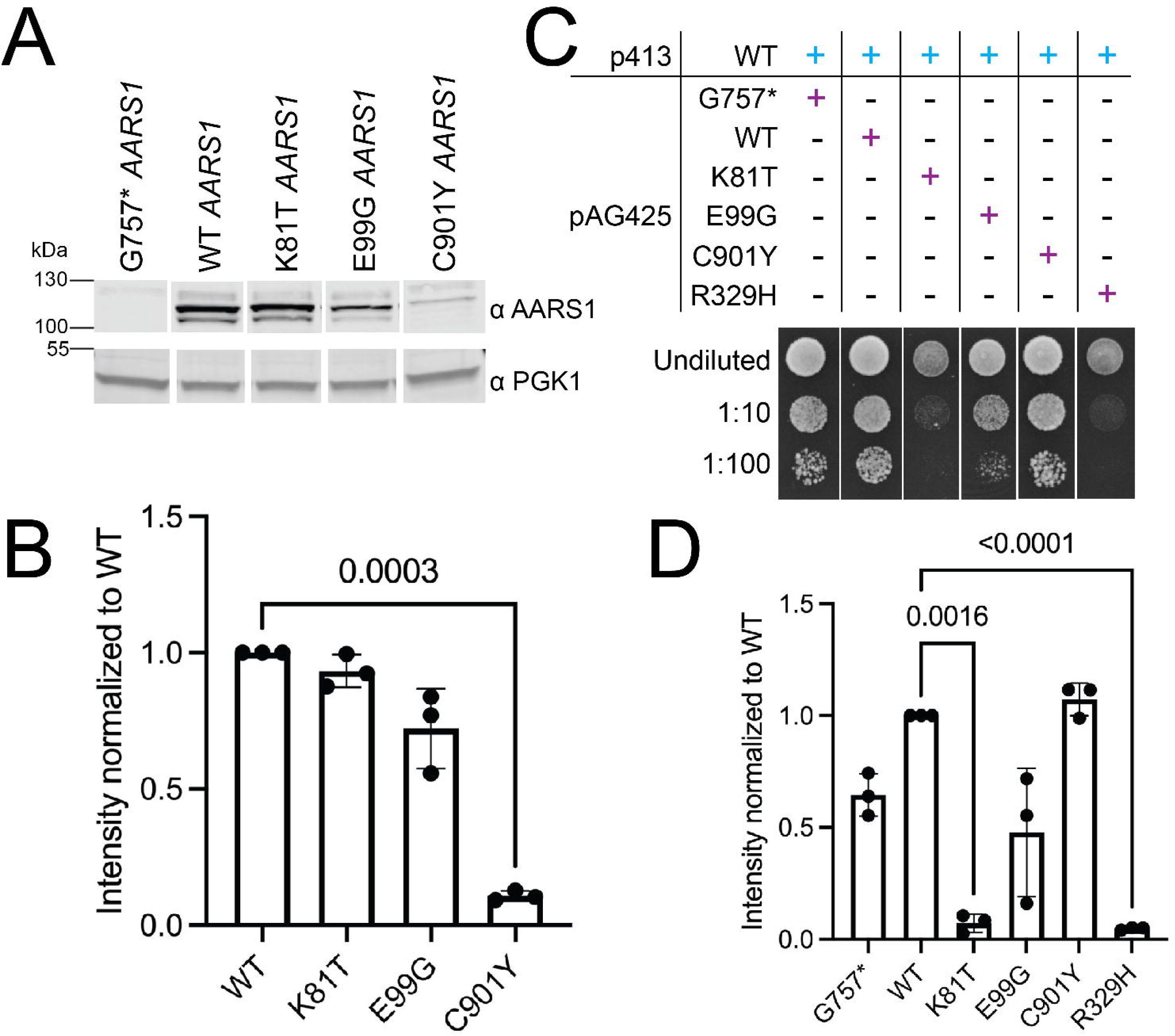
Protein expression and dominant-negative effects of loss-of-function *AARS1* missense variants. (**A**) AARS1 protein expression in transformed yeast. Western blot analyses were performed on protein lysates isolated from haploid yeast that were transformed with vectors containing the indicated inserts. The vector used in each experiment is across the top and sizes (kDa) are indicated at the left. (**B**) Band intensity was quantified relative to control PGK1 expression and normalized to wild-type AARS1 protein levels. (**C**) Haploid yeast with a doxycycline-repressible endogenous *ALA1* (the yeast ortholog of *AARS1*) and containing a wild-type (WT) *AARS1* p413 construct were transformed with pAG425 vectors containing the indicated *AARS1* allele. Resulting cultures were plated undiluted or diluted (1:10 or 1:100) on media containing doxycycline and grown at 30°C for five days. (**D**) Images of yeast spots were quantified to assess the relative growth of yeast co-expressing WT and *AARS1* mutants in comparison to yeast co-expressing WT *AARS1* on both p413 and pAG425. In both (B) and (D), the average intensity for three replicates was calculated and is depicted as the bar height. Error bars represent standard deviation. Statistical significance was determined by one-way ANOVA with the Geisser-Greenhouse correction and Dunnett’s multiple comparison’s test with individual variances computed for each comparison. Only comparisons that were statistically significant are annotated with a p-value.

### K81T AARS1 represses the function of wild-type AARS1 in yeast growth assays

K81T and E99G *AARS1* are loss-of-function alleles that do not impact protein levels. These data raise the possibility that these two alleles may have the potential to exert dominant-negative effects by interfering with the wild-type allele in the context of the AARS1 holoenzyme (a homodimer). We previously developed an assay to test for dominant-negative effects of *AARS1* variants by expressing mutant *AARS1* in the presence of wild-type *AARS1* and evaluating the effects on yeast cell growth (Meyer-Schuman et al., 2023). In brief, wild-type human *AARS1* on p413 and null (G757* *AARS1*), wild-type, or mutant human *AARS1* on pAG425 were transformed into the haploid ptetO7-*ALA1* yeast. Yeast growth was evaluated on medium containing doxycycline to repress endogenous *ALA1* expression and galactose to express the experimental allele; wild-type human *AARS1* was constitutively expressed on p413. When null *AARS1* on pAG425 was co-expressed with wild-type *AARS1* on p413, there was robust yeast cell growth, comparable to when wild-type *AARS1* on pAG425 is co-expressed with wild-type *AARS1* on p413 (**Figure 4B** and Supplementary Figure 4). This is consistent with wild-type human *AARS1* on p413 alone being sufficient to support yeast cell growth. When E99G or C901Y *AARS1* were co-expressed with wild-type *AARS1*, there was similar yeast cell growth compared to wild-type *AARS1* (**Figure 4B** and Supplementary Figure 4), indicating that these alleles do not impact the function of wild-type *AARS1*. In contrast, when K81T human *AARS1* was co-expressed with wild-type human *AARS1*, there was significantly reduced yeast cell growth compared to wild-type *AARS1* (p-value 0.0016; **Figure 4B** and Supplementary Figure 4). The reduction in yeast cell growth was similar to that associated with R329H *AARS1* (**Figure 4B** and Supplementary Figure 4), which causes dominant axonal neuropathy in several families (Meyer-Schuman et al., 2023). Importantly, the decreased yeast growth associated with co-expression of K81T human *AARS1* and wild-type human *AARS1* is rescued upon derepression of the endogenous *ALA1* yeast gene (Supplementary Figure 4B, middle column). This observation indicates that the reduced yeast growth is a direct effect of impaired alanyl-tRNA synthetase function and not due to off-target toxicity. In sum, our data suggest that K81T *AARS1*, a variant implicated in recessive disease, has dominant-negative properties and that it may cause axonal peripheral neuropathy in carriers of this variant.

## DISCUSSION

Here we describe a humanized yeast assay to comprehensively test and compare the functional consequences of 16 recessive-disease-associated *AARS1* variants using a humanized yeast model. First, variants were tested in a yeast complementation assay using a high-copy number vector. Six variants (K81T, E99G, S188H, V666A, T726A, and C901Y *AARS1*) displayed complete or partial loss-of-function. Next, the remaining variants that supported growth similar to wild-type *AARS1* were tested in a yeast complementation assay using a low-copy number vector to detect more subtle effects on function. Four additional variants (G581S, N604K, I699T, and G913D *AARS1*) displayed loss-of-function effects. Importantly, most of the missense variants tested (10 out of 16) demonstrated loss-of-function effects in our yeast system, consistent with previous studies and supporting partial loss of function as the molecular mechanisms of *AARS1*-associated recessive disease. Additionally, no patients carried two complete loss-of-function alleles, which is consistent with *AARS1* being essential for viability. Thus, yeast is an effective model for testing recessive *AARS1* alleles for loss-of-function effects. Data from this model will be useful toward building arguments for the pathogenicity of newly discovered *AARS1* alleles and similar studies should evaluate yeast as a model to study pathogenic alleles at other ARS loci.

That said, we noted two limitations in employing yeast to study *AARS1* alleles. First, this model was unable to detect loss-of-function effects for certain known pathogenic alleles. Two genotypes (Family B, R751G/R751G *AARS1*; and Family D, L298Q/R751G *AARS1*) consisted of two missense variants with neither variant demonstrating loss-of-function effects in one of our yeast assays; however, R751G is a highly confident pathogenic allele that has been identified in several patients and both R751G and L298Q *AARS1* were predicted to be likely pathogenic by AlphaMissense (**Table 2**) (Cheng et al., 2023). Furthermore, *in vitro* aminoacylation assays previously showed that R751G *AARS1* results in a 10-fold decrease in tRNA charging (**Table 2**) (Simons et al., 2015), and aminoacylation activity in L298Q/R751G *AARS1* patient fibroblasts was 37% compared to controls (Marten et al., 2020). Thus, while yeast is an effective system, there are limitations including that subtle differences in cell growth are difficult to observe and that yeast are single-celled organisms that may not demonstrate the functional consequences of variants in a complex, multicellular human (Oprescu et al., 2017). To address these limitations, orthogonal approaches (*e.g.*, enzyme kinetic assays) should be employed to assess for loss-of-function effects of *AARS1* alleles.

Second, our functional studies in yeast were unable to reveal relationships between *AARS1* gene dysfunction and clinical severity. The relationship between genotype and phenotype is challenging to evaluate given variability in clinical evaluations and difficulty in quantifying the severity of disease. Of the patients with bi-allelic missense variants, Family E (I699T/C901Y *AARS1*) had the most significant functional consequences in our assays with C901Y *AARS1* demonstrating loss of function when modeled in the high copy number vector and I699T *AARS1* showing reduced growth when modeled in the low copy number vector; however, the patient phenotype was restricted to non-photosensitive trichothiodystrophy (**Table 2**) (Botta et al., 2021). Helman et al. described a series of patients (Families G-P in **Table 2**) with either early infantile-onset, severe (Families G-J) or later-onset, milder (Families K-P) recessive disease, and they found no correlation between the amount of decreased enzyme activity in fibroblast lysate aminoacylation studies and age of onset or severity (Helman et al., 2021). Our yeast studies also do not reveal a relationship between loss-of-function effects in yeast complementation assays and disease severity for these patients (**Table 2**). In sum, while yeast is an effective model to study *AARS1* allele function, caution should be employed in interpreting negative data and in testing for phenotype-genotype correlations.

To identify potential dominant-negative *AARS1* alleles identified in patients with recessive disease, three complete loss-of-function variants (K81T, E99G, and C901Y *AARS1*) were assessed for effects on protein expression and for the ability to impact the function of wild-type *AARS1*. These efforts revealed that one allele (K81T) was expressed at levels comparable to the wild-type protein and resulted in reduced growth when co-expressed with wild-type *AARS1*, consistent with previously described dominant-negative alleles that cause axonal peripheral neuropathy (Meyer-Schuman et al., 2023). To our knowledge, no single *AARS1* variant has been implicated in both recessive and dominant diseases. Our data suggest that K81T *AARS1*, a variant implicated in recessive disease, could also result in dominant axonal neuropathy in heterozygous carriers (*i.e.*, parents and siblings of patients with the associated recessive phenotype). Furthermore, our results justify revisiting the clinical phenotype of the heterozygous parent in the affected pedigree (Simons et al., 2015) and suggest that *AARS1*-associated phenotypes exist along a spectrum where peripheral neuropathy is caused by loss-of-function, dominant-negative alleles that decreases *AARS1* with downstream effects on the integrated stress response (Spaulding et al., 2021), and that a multi-system syndrome is caused by two alleles with loss-of-function effects causing a further reduction in *AARS1* function. In summary, this study has important implications for studying the allelic and clinical heterogeneity, and the molecular mechanisms of *AARS1*-associated disease.

## Supporting information

Kuo supplementary materials

## ACKNOWLEDGEMENTS

A.A. is supported by a grant from the National Institute of General Medical Sciences (GM136441). M.E.K. is supported by the NIH Medical Scientist Training Program Training Grant (GM007863), the NIH Cellular and Molecular Biology Training Grant (GM007315), and an NIH National Research Service Award (F31) from the National Institute of Neurological Disorders and Stroke (NS113515).

