## Supplementary material for "Comprehensive assessment of recessive, pathogenic *AARS1* alleles in a humanized yeast model reveals loss-of-function and dominant-negative effects": Kuo supplementary materials

Supplementary Figure 1 displays all three yeast complementation assay replicates with all control plates for all 16 *AARS1* missense variant pAG425 constructs shown in Figure 2. Supplementary Figure 2 displays yeast complementation assay replicates and control plates for *AARS1* missense variants p413 constructs shown in Figure 3. Supplementary Figure 3 shows full blot replicates for western blot analyses in Figure 4A. Supplementary Figure 4 contains replicates and control plates for yeast complementation assays co-expressing loss-of-function variants with wild-type *AARS1* or empty vectors to test for dominant toxicity, a subset of which are shown in Figure 4B. Mutagenesis primer sequences for *AARS1* missense variants are in Supplementary Table 1.

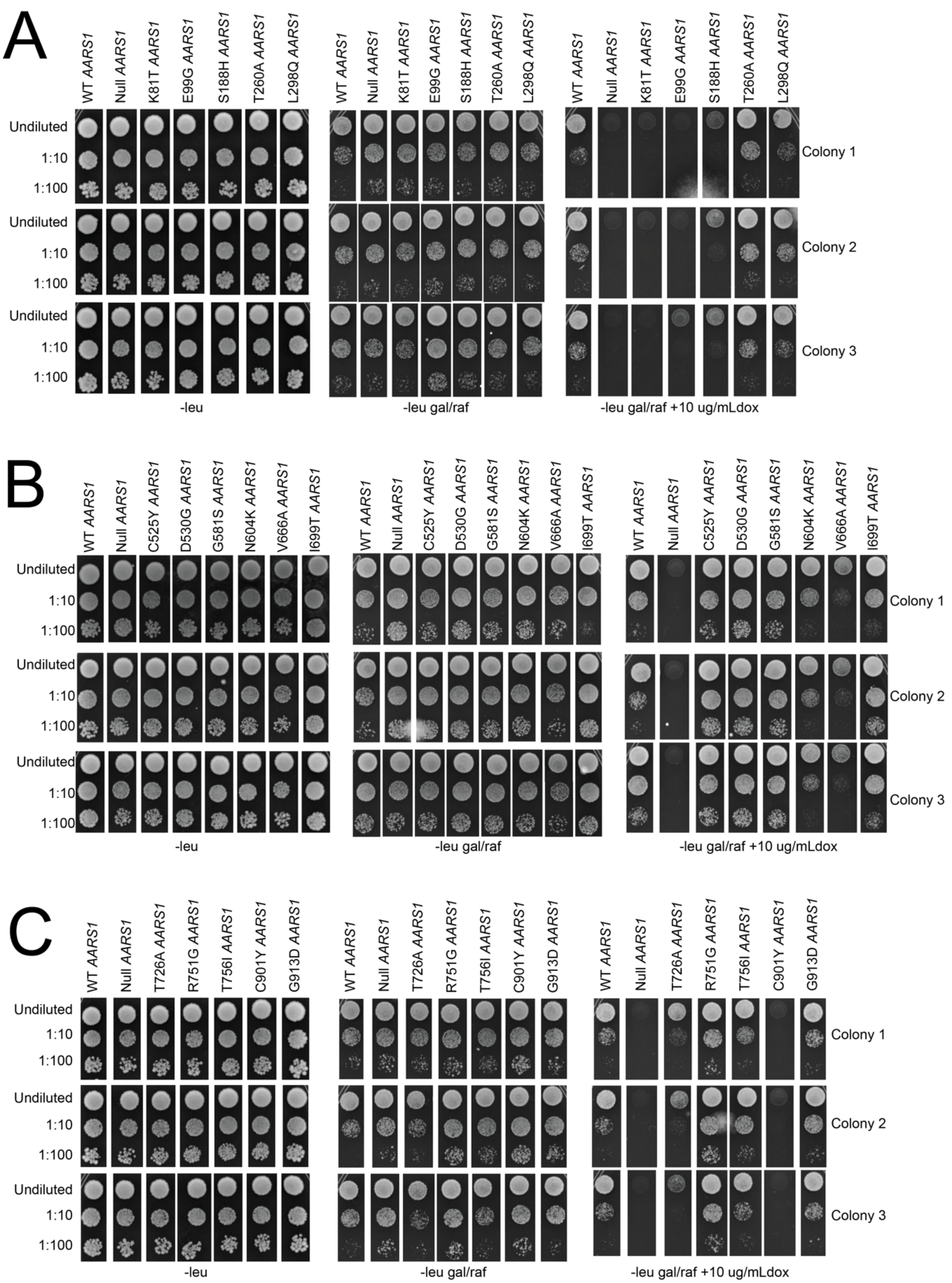


**Supplementary Figure 1**. Yeast complementation assay replicates and controls using a high-copy number vector (pAG425). Haploid yeast with a doxycycline-repressible endogenous *ALA1* (the yeast ortholog of *AARS1*) were transformed with pAG425 vectors containing wild-type (WT) *AARS1* or mutant *AARS1*, or a vector with a null allele (G757* *AARS1*); the vector used in each experiment is indicated across the top. Resulting cultures were plated undiluted or diluted (1:10 or 1:100) on glucose media lacking leucine, galactose/raffinose media lacking leucine, or galactose/raffinose media lacking leucine and containing doxycycline. Yeast were grown at 30°C for five days. Variants were divided into three sets (**A**, **B**, and **C**). Bacterial contamination was present on some plates but did not interfere with interpretation or quantification.

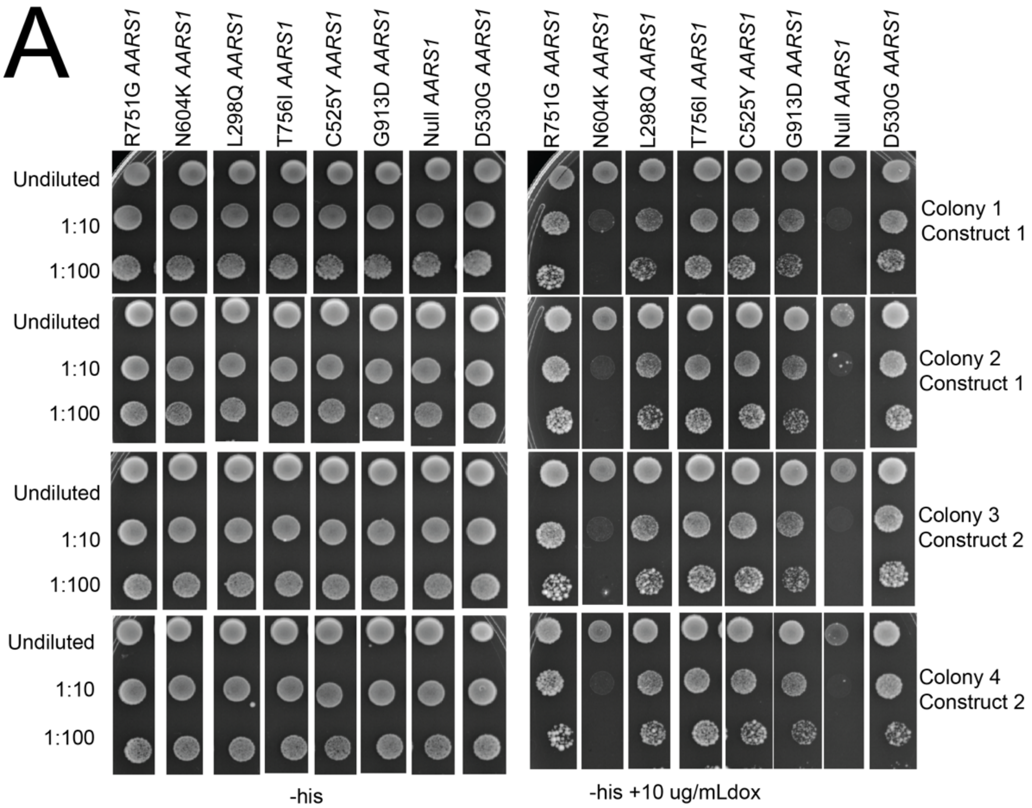


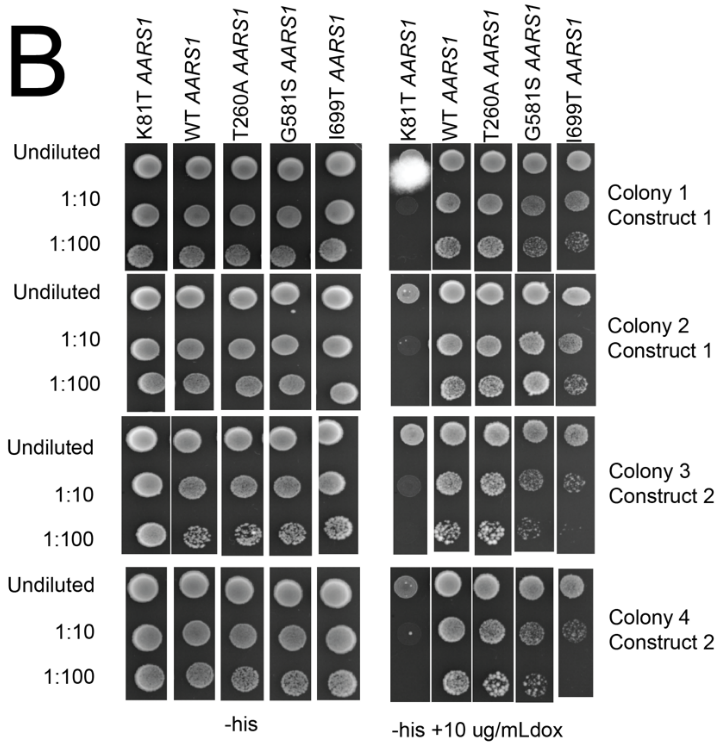


**Supplementary Figure 2**. Yeast complementation assay replicates and controls using a low-copy number vector (p413). Haploid yeast with a doxycycline-repressible endogenous *ALA1* (the yeast ortholog of *AARS1*) were transformed with p413 vectors containing wild-type (WT) *AARS1* or mutant *AARS1*, or a p413 vector with a null allele (G757* *AARS1*); the vector used in each experiment is indicated across the top. Resulting cultures were plated undiluted or diluted (1:10 or 1:100) on glucose media lacking histidine or glucose media lacking histidine and containing doxycycline. Yeast were grown at 30°C for five days. Variants were spotted across two plates (**A** and **B**). Bacterial contamination was present on some plates but did not interfere with interpretation or quantification.

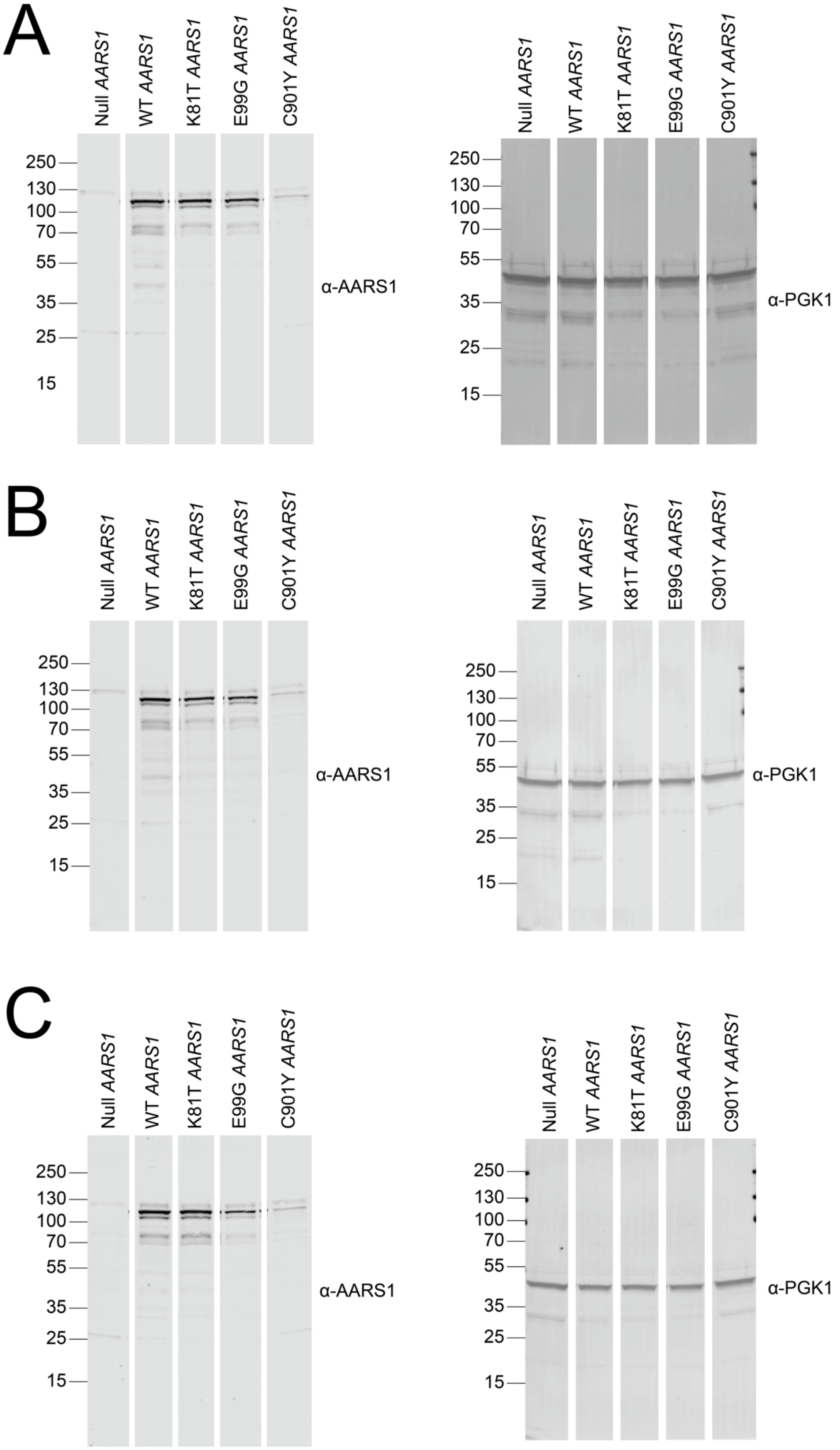


**Supplementary Figure 3**. Three replicates (**A**, **B**, and **C**) and full blots of western blot experiments. Western blot analyses were performed using protein lysates isolated from haploid yeast that were transformed with the pAG425 vector indicated across the top and antibodies to AARS1 and PGK1.

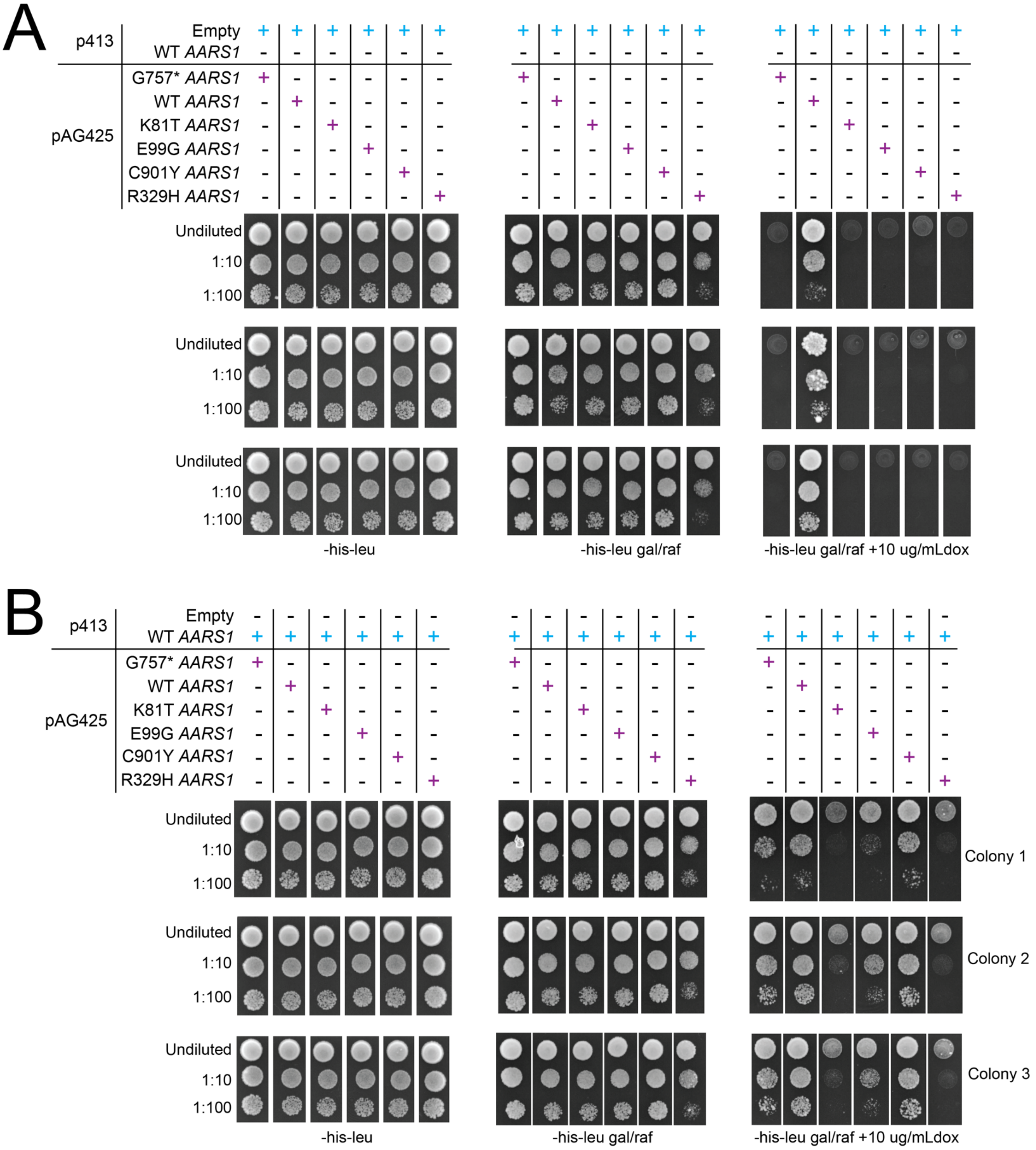


**Supplementary Figure 4**. Replicates and control plates for yeast complementation assays co-expressing 'empty' (*i.e.*, with no *AARS1* insert) (**A**) or wild-type *AARS1* p413 (**B**) constructs with mutant *AARS1* pAG425 constructs. Haploid yeast with a doxycycline-repressible endogenous *ALA1* (the yeast ortholog of *AARS1*) containing a wild-type (WT) *AARS1* or 'empty' p413 construct were transformed with pAG425 vectors containing the indicated insert. Resulting cultures were plated undiluted or diluted (1:10 or 1:100) on glucose media lacking histidine and leucine, galactose/raffinose media lacking histidine and leucine, or galactose/raffinose media lacking histidine and leucine and containing doxycycline. Yeast were grown at 30°C for five days.
